# Human plasma-like medium improves T lymphocyte activation

**DOI:** 10.1101/740845

**Authors:** Michael A. Leney-Greene, Arun K. Boddapati, Helen C. Su, Jason Cantor, Michael J. Lenardo

## Abstract

T lymphocytes are critical for effective immunity and the ability to study their behavior in synthetic media *in vitro* facilitates major discoveries in their development, function, and fate. However, the composition of human plasma differs from synthetic media and we hypothesized that these differences could have important effects on cell physiology. We therefore compared T lymphocyte activation in human plasma-like medium (HPLM) to RPMI supplemented with dialyzed FBS (RPMI^dFBS^) and found that it entrained markedly different transcriptional responses. We also found that the concentration of calcium in RPMI^dFBS^ is six-fold lower than HPLM causing altered T cell activation which could be reversed by calcium addition. Thus, investigators should be cognizant of differences between commonly used media formulations and HPLM which is based on the *in vivo* plasma environment as these could profoundly affect their experimental results. Physiologic media may be a valuable new way to study immune cells in culture.

## INTRODUCTION

Adaptive immune systems involve the movement of immune cells throughout the internal milieu of the blood and interstitial spaces to recognize foreign agents and respond, usually through transcriptional changes, to ensure the health of the host. T lymphocytes are central effectors of immunity in infectious diseases, autoimmunity, and cancer and their metabolic status is critical for shaping an appropriate T cell response. Recent publications have highlighted the influence of lipids, glucose, and amino acid metabolism on both the magnitude and characteristics of T cell responses (Pearce *et al.*, 2009; van der Windt and Pearce, 2012; Sinclair *et al.*, 2013; O’Sullivan *et al.*, 2014; Buck *et al.*, 2017; Ma *et al.*, 2017; Werner *et al.*, 2017; Jacobs *et al.*, 2018).

Formulations of common cell culture media such as RPMI-1640 were developed in the mid-20^th^ century to optimize *in vitro* growth of cell lines and have since undergone remarkably little change (Eagle, 1955; McCoy, Maxwell and Kruse, 1959; Moore *et al.*, 1966). Despite the new focus on the effects of metabolic changes during T cell activation and proliferation, culture conditions that more closely resemble the *in vivo* milieu have not been studied. Recently, several studies in non-immune cells have described the use of modified traditional media or new systematically constructed synthetic media with the aim of either improving growth in cell culture or better modeling the in vivo environment (Favaro *et al.*, 2012; Schug *et al.*, 2015; Pan *et al.*, 2016; Cantor *et al.*, 2017; Voorde *et al.*, 2019). Among these is Human Plasma-Like Medium (HPLM), which contains a cocktail of 31 components that are absent from the defined formulations of RPMI and other standard basal culture media (Cantor *et al.*, 2017). Furthermore, HPLM contains physiological concentrations of other common media components such as glucose, amino acids, and salt ions. All of these components may be present at non-physiological levels in FBS, the most widely used tissue culture supplement (Cantor *et al.*, 2017). Here, we asked whether HPLM altered the *in vitro* activation of primary human T lymphocytes compared to RPMI.

## RESULTS

### Transcriptome analysis reveals extensive differences in T lymphocytes activated in HPLM compared to RPMI^dFBS^

T lymphocytes *in vivo* undergo broad transcriptional re-programming following TCR activation. *In vivo* this occurs in a rich internal milieu containing high levels of amino acids, lipids, and a variety of small organic metabolites, whereas typical *in vitro* culture methods using RPMI only contain a skeleton of essential amino acids, vitamins, glucose, and salts (Eagle, 1955; McCoy, Maxwell and Kruse, 1959; Crabtree, 1989; Cantor *et al.*, 2017). We therefore tested activation in HPLM compared to RPMI^dFBS^ during T cell activation (Cantor *et al.*, 2017) (Figure 1A, Supplemental Table 1). We supplemented both RPMI and HPLM with dialyzed (hereafter referred to as RPMI^dFBS^ and HPLM, respectively) rather than complete FBS (RPMI^cFBS^) in order to control the levels of metabolites and ions in both preparations, while still providing protein growth factors required for cell survival. We then activated purified human naïve T lymphocytes from three individual donors with plate-bound anti-CD3/CD28 antibodies for 48 or 120 hours in either HPLM or RPMI^dFBS^, isolated polyadenylated mRNAs and characterized the transcriptional differences between these two conditions via deep sequencing. Principal component analysis revealed changes between 48 and 120 hours of activation independently of the medium used. Nonetheless, the 2^nd^ and 3^rd^ principal components divided each group of samples (RPMI^dFBS^-48 hours, RPMI^dFBS^-120 hours, HPLM-48 hours and HPLM-120 hours) into clear clusters revealing the transcriptional differences between HPLM and RPMI^dFBS^ (Figure 1C). We next used gene set enrichment analysis (GSEA) to identify statistically significant differences in 29 different Kyoto encyclopedia of genes and genomes (KEGG) pathways (Kanehisa and Goto, 2000; Kanehisa *et al.*, 2019). Nine pathways were significantly different at 48 hours, 19 significantly different at 120 hours and one pathway was shared between both timepoints (Supplemental Figure 1). Among these we observed a striking enrichment of pathways involved in DNA replication and cell cycle in HPLM at 120 hours post-activation and an enrichment of pathways involved in T cell activation at 48 hours (Supplemental Figure 1). In particular, essentially every gene in the KEGG DNA Replication pathway exhibited increased expression in HPLM relative to RPMI^dFBS^ (Figure 1D). Thus, T cell activation in HPLM was superior to RPMI^dFBS^ and this difference was readily apparent as early as 48 hours post activation.

**Figure 1.**
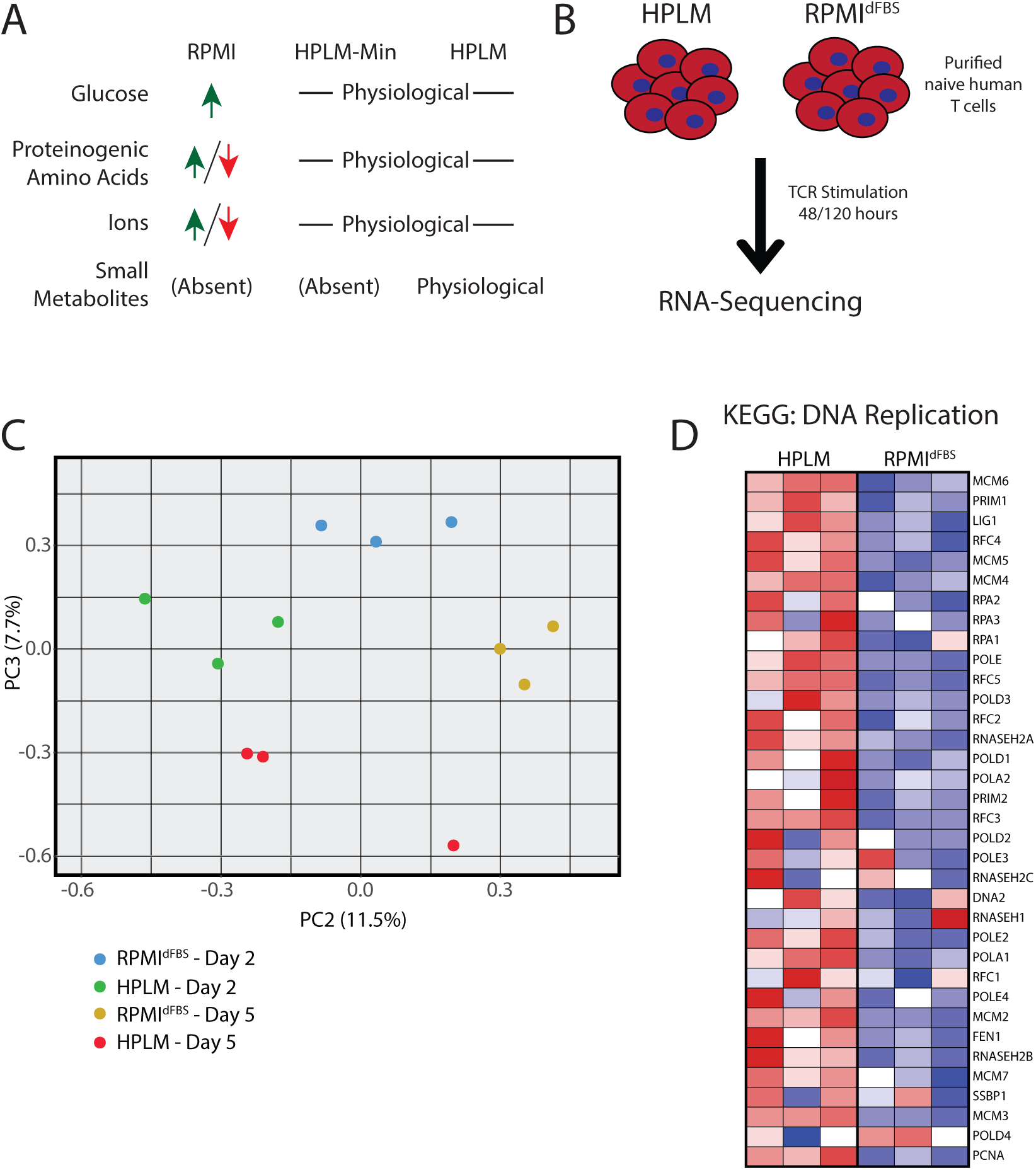
Transcriptomic analysis reveals major differences between lymphocytes activated in HPLM compared to RPMI. (A) Schematic of the comparative compositions of RPMI, HPLM and HPLM-Min. Detailed composition can be found in Supplemental Table 1. (B) Experimental outline for T cell activation in either HPLM or RPMI and downstream transcriptome analysis. (C) PCA plots showing principal components 1 and 2 in primary naïve human T cells activated for 48/120 hours in either HPLM or RPMI. (D) Heatmap showing the Log_2_(fold change) in transcript abundance for genes involved in the KEGG DNA Replication pathway.

Metabolic pathways also showed important transcriptional differences between the media. Transcriptional differences of some of the rate-limiting enzymes for each of these pathways are highlighted in Table 1. We found major increases in genes involved in amino acid metabolism in HPLM, including the KEGG pathways arginine/proline metabolism, glycine/threonine/serine metabolism and alanine, aspartate and glutamate metabolism. These distinctions are most likely being driven by the roughly 2 to 10 fold excesses of arginine, aspartate, serine and glutamate present in RPMI relative to HPLM (Supplemental Table 1), and thus human plasma which might suppress their anabolic pathways. Also notable were increases in nucleic acid metabolism including DNA repair, pyrimidine metabolism, RNA polymerase, splicosome, and homologous recombination. Finally, the p53 signaling pathwas also increased in HPLM. Conversely, genes involved in glycerophospholipid and cytochrome p450 metabolism as well as cell adhesion, allograft rejection, graft vs. host disease, type I diabetes, cytokine receptor interactions, and autoimmune thyroid disease were significantly decreased in HPLM. Together these data suggest a more favorable T cell response to promote proliferation and restrain autoimmunity.

**Table 1:** Transcriptional differences in metabolic genes in T cells cultured in HPLM.

| **Serine biosynthesis** |  |
| --- | --- |
|  | **Fold change in expression in HPLM** |
| **PHGDH** | 6.10 |
| **PSAT1** | 5.30 |
| **PSPH** | 1.25 |
| **Glycine, Arginine, Proline biosynthesis** |  |
| **SHMT1** | 1.80 |
| **SHMT2** | 1.33 |
| **PYCR1** | 4.32 |
| **ASS1** | 81.00 |
| **Aspartate biosynthesis** |  |
| **GOT1** | 1.45 |
| **Cholesterol biosynthesis** |  |
| **HMGCS1** | 0.38 |
| **HMGCR** | 0.49 |
| **MVD** | 0.48 |
| **FDFT1** | 0.47 |

### HPLM is superior to RPMI^dFBS^ in supporting naive human T cell activation

Guided by our RNA-sequencing data, we hypothesized that T cell activation and proliferation would be more efficient in HPLM. We therefore measured markers of activation in naïve human T cells stimulated as in Figure 1. In both CD4^+^ and CD8^+^ T cells from five different healthy donors, we observed a significant increase in CD25 and CD69 in HPLM compared to RPMI^dFBS^ in (Figure 2A). Higher concentrations of anti-CD3/CD28 antibodies gave nearly equivalent maximal response rates in both RPMI^dFBS^ and HPLM (*i.e.* >90% positive for both CD25 and CD69) (Figure 2A). This suggests that t HPLM lowers the activation threshold. This was visually apparent as there were large clusters of activated T cells in the HPLM cultures that were absent from cells cultured in RPMI^dFBS^ as well as a significant increase in cell size measured by flow cytometry (Figure 2B). Finally, as we anticipated from the transcriptome changes, with the lower dose of TCR stimulation, we also observed vigorous proliferation of cells in HPLM that was hardly apparent in RPMI^dFBS^ (Figure 2C).

**Figure 2.**
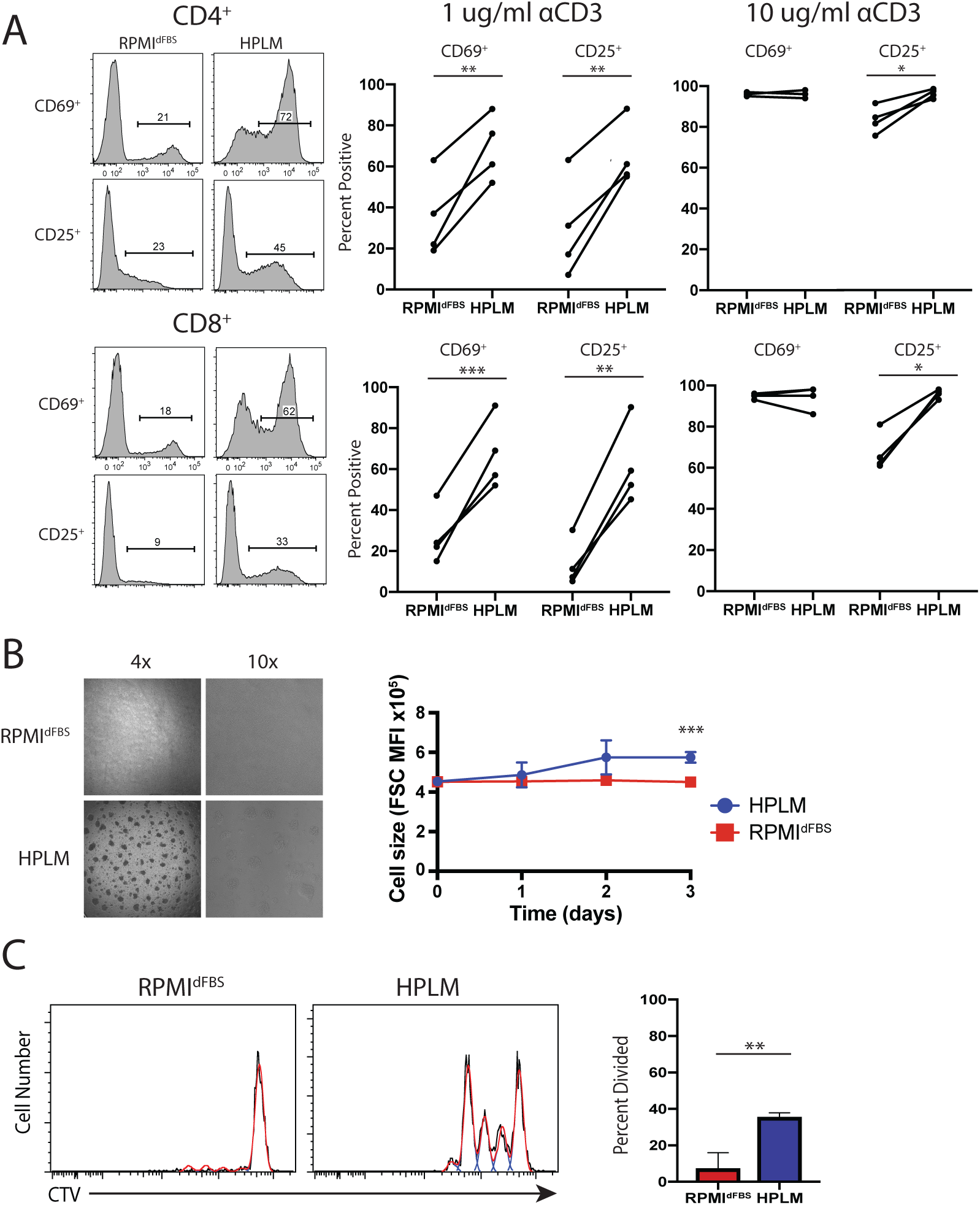
Physiologic media is superior to RPMI for human T cell activation. (A) Flow cytometric measurement of T cell activation markers CD25 and CD69 on purified naïve human T cells following stimulation with either 1 or 10 μg of plate-bound anti-CD3 (αCD3) /CD28 in HPLM or RPMI. Data shown are representative of 4 different experiments each conducted with cells isolated from a different healthy donor (Student’s t test; paired; two-tails; *p < 0.05, **p < 0.01, ***p < 0.001). (B) Bright-field microscopy images of naïve T cells stimulated with 1 μg of plate-bound anti-CD3/CD28 in either HPLM or RPMI. (C) Flow cytometry histograms of Celltrace Violet staining of naïve T cells stimulated in either HPLM or RPMI (left) with quantitation of the fraction of CD4^+^ T cells that have undergone at least one division (right). Columns represent the mean of three experiments, each done with cells isolated from a different healthy donor (Student’s t test; unpaired; two-tails; *p < 0.05).

### Physiological calcium availability augments T cell activation

We next examined which components of HPLM improve T cell activation. We compared the activation of T cells in HPLM, HPLM-Min (which lacks the 31 small metabolites but has the same concentration of amino acids, glucose and ions, see Supplemental table 1 and previous publications (Cantor *et al.*, 2017)) and RPMI^dFBS^. The broad compositions of these three media are depicted in Figure 1A while the precise concentration of each substituent is given in Supplemental Table 1. We also included a condition in which we added all HPLM specific polar metabolites back to RPMI as well as dialyzed FBS (RPMI-Metabolites). We observed equivalent activation in the HPLM and HPLM-Min conditions, and significantly decreased activation in either RPMI^dFBS^ and RPMI-Metabolites (Figure 3A). These results suggest that the polar metabolites in HPLM are dispensable for T cell activation and rather concentrations of amino acids, glucose or ions are responsible. One striking difference in composition between HPLM and RPMI^dFBS^ was the increased Ca^2+^ concentration [Ca^2+^] in HPLM (2.4 mM) compared to RPMI (0.4 mM) (Supplemental Table 1), which is markedly hypocalcemic relative to the *in vivo* milieu (2-2.5 mM) (Goldstein, 1990). Indeed, supplementation of RPMI^dFBS^ with 2 mM CaCl_2_ (RPMI+Ca^2+^) was sufficient to fully activate cells as well as in HPLM (Figure 3A).

**Figure 3.**
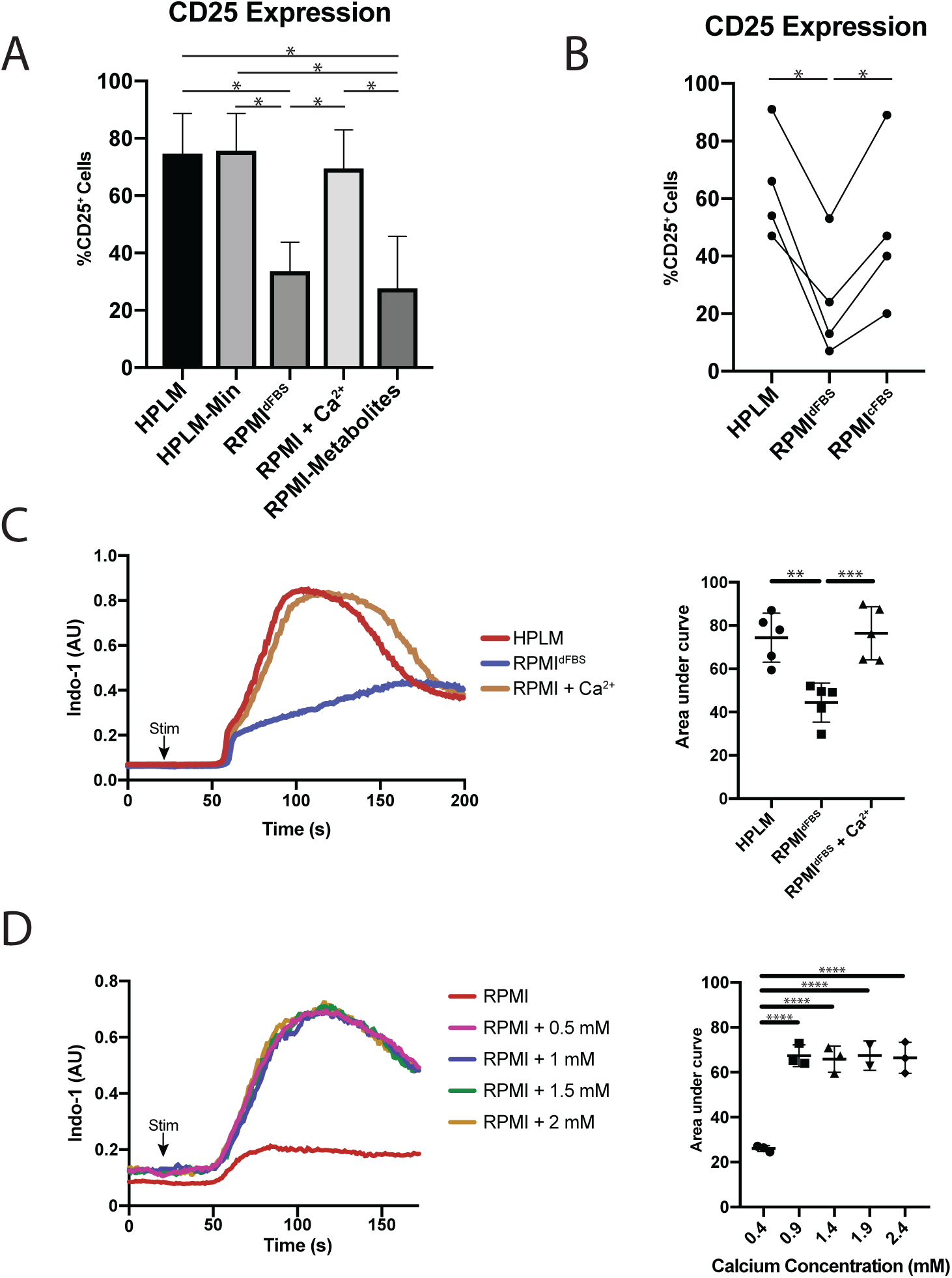
RPMI is Severely Hypocalcemic Relative to Human Plasma. (A) Measurement of activation marker CD25 on CD4^+^ T cells comparing RPMI and minimal-HPLM supplemented with various metabolite components unique to HPLM. Columns represent the mean with error bars showing the standard error (One-way ANOVA comparing other conditions to HPLM with p values calculated by Dunnett test; *p < 0.05 (B) Measurement of activation marker CD25 on CD4^+^ T cells activated in HPLM, RPMI^dFBS^ or RPMI^cFBS^. Experiment was repeated four times with each repeat using a different healthy human donor (One-way ANOVA; Tukey’s test; *p < 0.05). (C) Flow cytometric plots of calcium flux following primary stimulation of isolated human CD8^+^ T lymphocytes in either RPMI or HPLM (left) as well as quantification of the area under the curve (right). Quantification shows data from five experiments each done with a different individual donor and error bars show standard error (One-way ANOVA; Tukey’s test; **p < 0.01, ***p < 0.001). (D) Flow cytometric plots of calcium flux following primary stimulation of isolated human CD8^+^ T lymphocytes in RPMI supplemented with the indicated concentrations of calcium chloride. Quantification shows data from five experiments each done with a different individual donor and error bars show standard error (One-way ANOVA; Tukey’s test; **p < 0.01, ***p < 0.001).

Basal media used to culture T lymphocytes (typically RPMI) are traditionally supplemented with 10% unmodified FBS, and thus we next asked how such supplementation affects the Ca^2+^ levels of RPMI and T lymphocyte activation. We found that [Ca^2+^] = 3.9 mM in our typical FBS, and thus, a 10% supplement would result in a final [Ca^2+^] of ∼ 0.8 mM in RPMI^cFBS^. This value is still hypocalcemic but closer to a physiological [Ca^2+^]. We observed that TCR stimulation of naïve T cells in RPMI^cFBS^ produced nearly equivalent activation as in HPLM (Figure 3B). This suggests that increase [Ca^2+^] achieved by adding non-dialyzed FBS is sufficient to lead to normal activation. We next examined Ca^2+^ flux following TCR stimulation in either RPMI^dFBS^, RPMI^Ca2+^ and HPLM. It is worth noting that most calcium flux protocols are carried out in Hank’s balanced salt solution or Ringer’s solution, however some studies have shown decreased calcium flux in RPMI relative to these media (Prakriya *et al.*, 2006; Gwack *et al.*, 2008; Bertin *et al.*, 2014). We observed a striking decrease in the amount of calcium entering the cells following activation in RPMI^dFBS^ when compared to either HPLM or RPMICa^2+^ which were nearly equivalent (Figure 3C). Thus, the high activation threshold in RPMI was due to reduced Ca^2+^ in the medium.

Our results raised the possibility that calcium levels *in vivo* could influence T cell activation, and that hypocalcemic/hypercalcemic patients could be more prone to immunodeficiency or autoimmunity respectively. Previously it was shown that the extracellular [Ca^2+^] can influence cytokine production in murine T cells (Zimmermann, Radbruch and Chang, 2015). Moreover, our observations indicated that RPMI^dFBS^ poorly recapitulated *in vivo* conditions because the [Ca^2+^] was too low and could introduce variation in results depending on uncontrolled levels of Ca^2+^ in FBS. To test these ideas, we titrated calcium levels in basal (FBS free) RPMI in 0.5 mM increments up to a maximum of 2.2 mM and then measured TCR - induced calcium flux. We again observed that basal RPMI gave a meager Ca^2+^ flux, but [Ca^2+^] from 0.7 to 2.2 mM yielded virtually identical magnitude and kinetics of calcium fluxes (Figure 3D). This suggests that at the lowest physiological extracellular [Ca^2+^], the rate of Ca^2+^ flux can reach a maximum value and higher extracellular [Ca^2+^] likely do not affect the sensitivity of T cells to activation *in vivo*. However, the [Ca^2+^] in RPMI is insufficient to promote full Ca^2+^ flux or T cell activation without an additional source of Ca^2+^.

### HPLM promotes CD8^+^ T cell effector cytokine production

We next evaluated cytokine production in CD8^+^ T cells activated in HPLM or HPLM-Min. We activated T cells in RPMI^dFBS^ and grew them for 14 to 21 days supplemented with IL-2 prior to restimulation and measurement of cytokine production. This showed that TNFα and IFNγ production was equivalent in HPLM or HPLM-Min but substantially greater than RPMI^dFBS^ (Figure 4A), the same results were obtained with phorbol myristate acetate (PMA) and ionomycin or anti-CD3 antibodies crosslinked with protein A. Th IFNγ production was reduced in HPLM compared to HPLM-min though this did not reach statistical significance. We next examined whether this difference was due to the media composition during the initial stimulation or the subsequent restimulation by activating and expanding the cells in HPLM or RPMI^dFBS^ and then switching them to other media just prior to restimulation (Figure 4B). W Under these conditions, we observed robust cytokine production in cells that were activated in HPLM or RPMI^dFBS^ and then transferred to HPLM or RPMI^dFBS^ supplemented with 2mM Ca^2+^ (Figure 4B). However, irrespective of starting medium, those transferred to RPMI^dFBS^ with no calcium supplement showed a poor response indicating that having optimal medium, presumably optimal extracellular [Ca^2+^], is necessary at the time of stimulation.

**Figure 4.**
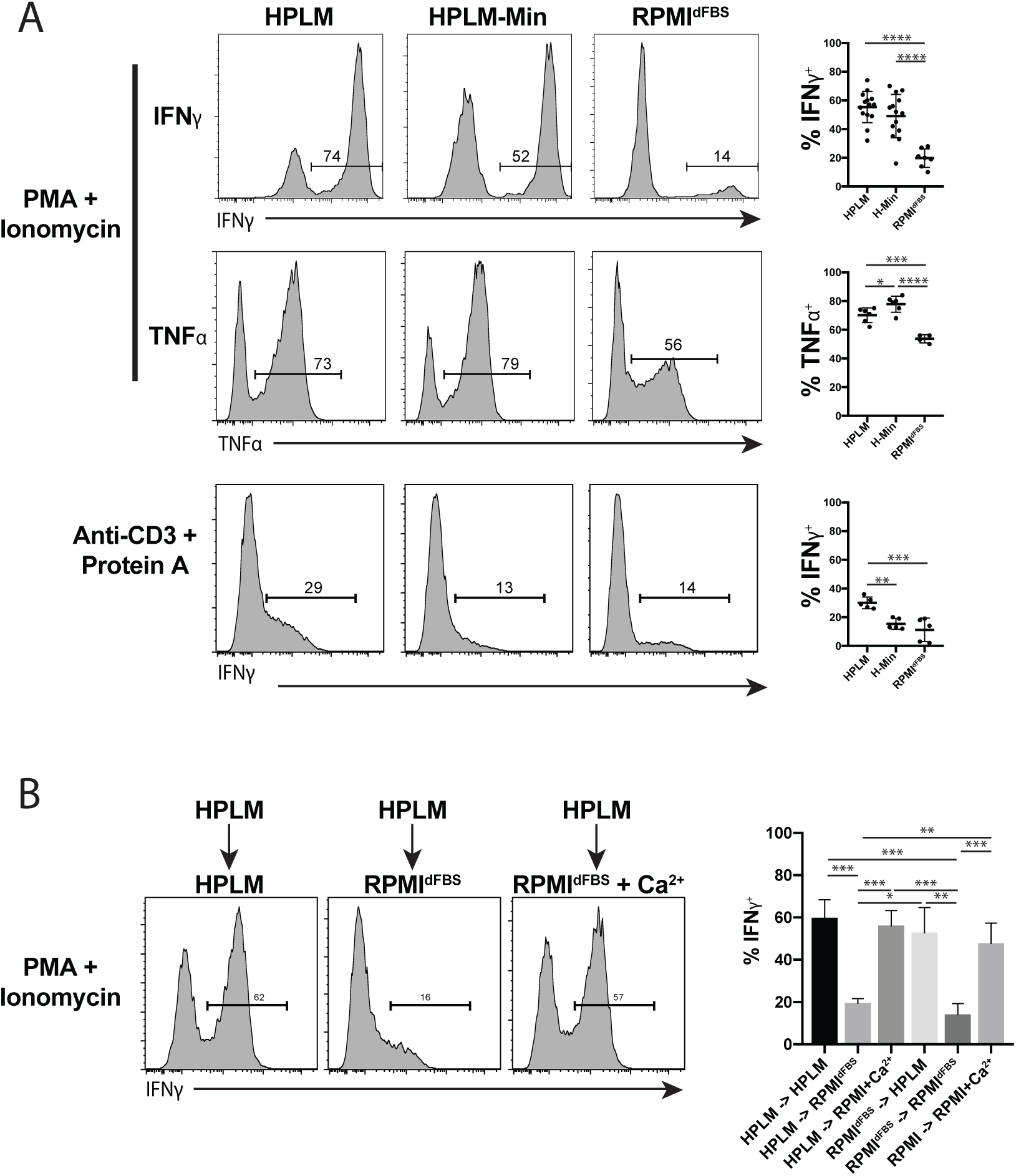
Activation in Physiologic Media leads to Superior Cytokine Production. (A) Levels of cytokines produced in primary human T lymphocytes following restimulation after being expanded in the indicated medium. T lymphocytes from 5 to 14 individuals were tested across 2 to 3 experiments, with the error bars representing standard deviation (One-way ANOVA; Tukey’s test; *p < 0.05, **p < 0.01, ***p < 0.001, **** p < 0.0001). (B) Quantification of cytokine production in primary human T lymphocytes expanded in the indicated medium, switched to fresh medium of the indicated type and then restimulated. Columns represent the mean of measurements from 3 experiments each with two individuals, with the error bars representing the standard error (One-way ANOVA; Tukey’s test; *p < 0.05, **p < 0.01, ***p < 0.001).

### CD19 CAR-T cell transduction efficiency in HPLM is similar to other commonly used media

T lymphocytes have become a central focus in the field of cancer immunotherapy, and many clinical protocols require the *in vitro* culture and expansion of human T cells engineered with chimeric antigen receptors (CAR-T cell) (Newick *et al.*, 2018). We therefore hypothesized that T cells cultured in HPLM might exhibit improved transduction efficiency with lentiviruses expressing CAR-T cell receptors. We compared HPLM to two other synthetic media typically used in clinical protocols: a mixture of AIM V and RPMI (supplemented with 5% human serum, referred to as AIM V here) and X-VIVO 15 (serum free). We observed equivalent CD25 expression in HPLM, HPLM-Min and AIM V media but much less with X-VIVO 15 after stimulation with plate-bound anti-CD3/CD28 antibodies (Figure 5A). Moreover, lentiviral transduction efficiency of T cells of both CD4^+^ and CD8^+^ T cells was equivalent in HPLM, HPLM-Min and AIM V media but was less with X-VIVO 15 though this did not reach statistical significance (Figure 5B). Taken together, HPLM also performs comparably or better than commonly used culture media for the generation of CAR-expressing T cells.

**Figure 5.**
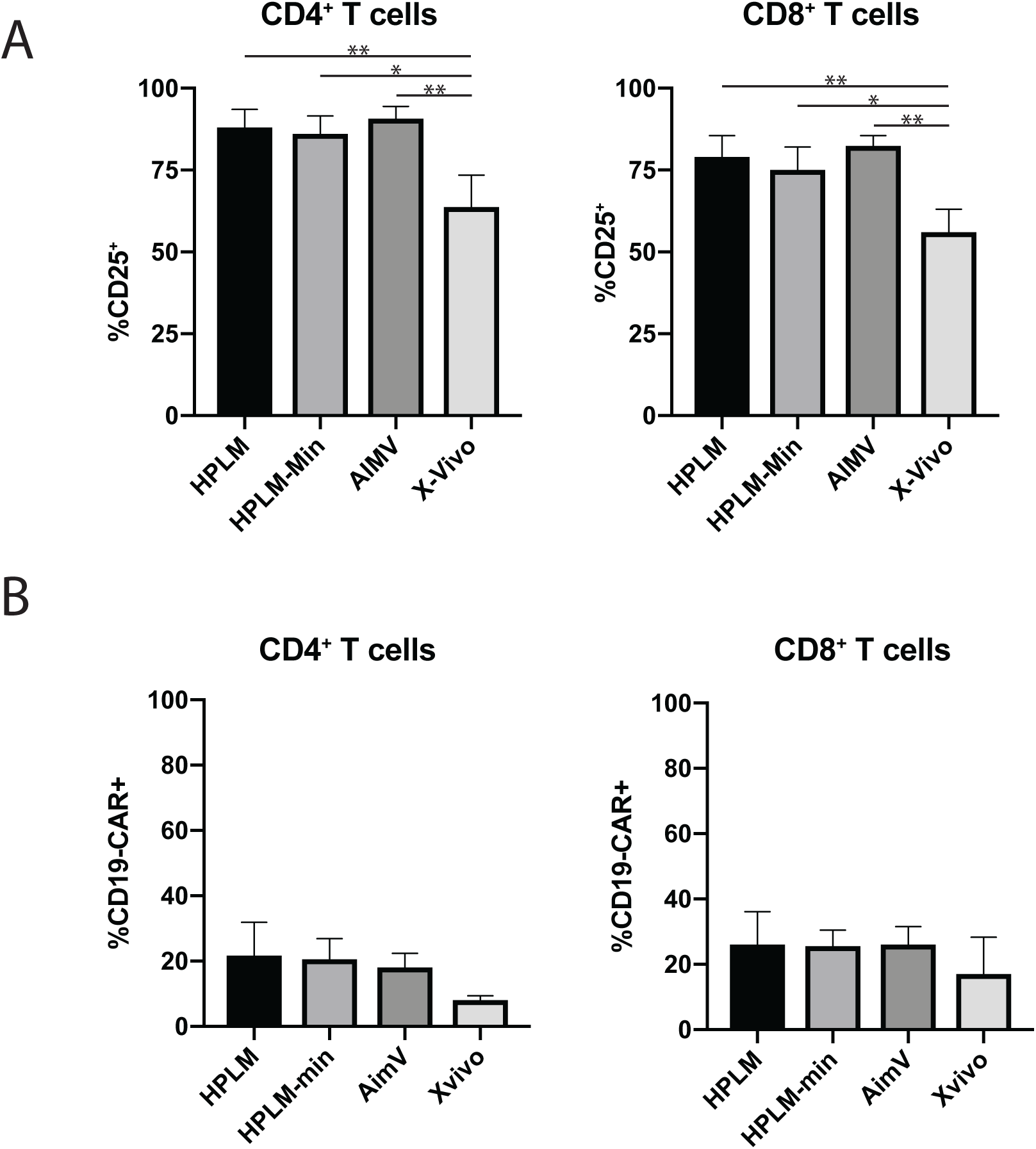
Similar Transduction Rates in HPLM and RPMI. (A) Quantification of levels of activation markers in CD4^+^ (left panel) and CD8^+^ T cells following activation in the indicated medium. Columns represent the mean of measurements from 3 experiments each conducted with cells from a different individual, with error bars representing the standard error (One-way ANOVA; Tukey’s test; *p < 0.05, **p < 0.01). (B) Quantification of transduction efficiency human CD4^+^ (left panel) and CD8^+^ T cells (right panel) with CD19-CAR expressing lentiviruses activated in the indicated media. Columns represent the mean of measurements from 3 experiments each conducted with cells from a different individual, with error bars representing the standard error (One-way ANOVA; Tukey’s test).

## DISCUSSION

The development of cell culture techniques in the mid 20^th^ century heralded an enormous advance in life sciences research by permitting tissue-free studies *in vitro* of cell physiology. However, media formulations developed in the 1960s are still universally used today. In this study, we extended a recent effort to modernize the field using the development of a cell culture medium modeled closely after the *in vivo* plasma substituents to the study of primary human lymphocytes. Our current study showed much more robust activation as well as secretion of effector cytokines by naïve human T cells *in vitro* in HPLM compared to RPMI^dFBS^. Furthermore, RNA-sequencing revealed a large number of underlying metabolic differences between culture in HPLM vs. RPMI^dFBS^ which could be important for investigators who are studying T cell metabolism. We also found that calcium is rate-limiting for lymphocyte activation in RPMI^dFBS^, which fits with recent studies demonstrating the molecular machinery involved in the acute Ca^2+^ flux that occurs after TCR engagement (Zhang *et al.*, 2005; Prakriya *et al.*, 2006). We found that RPMI^dFBS^ contains [Ca^2+^] that is insufficient to support full T cell activation. Other groups have observed increased *in vitro* secretion of effector cytokines in IMDM ([Ca^2+^] = 1.5 mM) or calcium supplemented RPMI^dFBS^ relative to basal RPMI in murine T cells *ex vivo* (Zimmermann, Radbruch and Chang, 2015). We also find that activation, proliferation and effector cytokine secretion in primary human T cells is reduced in RPMI^dFBS^ without Ca^2+^ supplementation. It is likely that the use of FBS as a common supplement to cell culture augments the calcium concentration of basal RPMI sufficiently, as we measured our current stock of complete FBS at 3.9 mM – raising the concentration to roughly 0.8 mM when added at a concentration of 10%. Although our current study suggests that this is enough to achieve maximal TCR -induced Ca^2+^ flux and activation, it is now clear that this is an independent variable that should be considered. Moreover, even RPMI^cFBS^ still corresponds to a severely hypocalcemic condition for T cell studies. For example, it is possible that the cross-linking anti-CD3 antibody used in the calcium flux masks an underlying defect that would be present in response to physiological antigens. Furthermore, the calcium flux assay only measures calcium changes for the first several minutes of a response, whereas the activation process is much more dynamic and can extend over the course of hours/days *in vivo*.

Looking beyond calcium, a comparison of great interest to us was the HPLM versus HPLM-min conditions to understand the impact of the plasma metabolites on T cell activation and proliferation. We found that these two media revealed only subtle differences in CD25/CD69 expression, proliferation or cytokine production after stimulation. However, the major differences observed in HPLM/HPLM-min and RPMI^dFBS^, suggests that the concentrations of extracellular calcium and amino acids play a larger role than the small plasma metabolites in T cell activation. One caveat is that previous work using HPLM used a much more robust dialysis procedure for production of the dialyzed FBS (Cantor *et al.*, 2017). It is possible that our commercial source was less stringently dialyzed and thus was still contaminated with appreciable levels of polar metabolites, masking any potential differences between T cells activated in HPLM and HPLM-min.

Comparison of the transcriptomes of T lymphocytes activated in HPLM versus RPMI^dFBS^ initially revealed major differences in activation which led to our subsequent focus on extracellular [Ca^2+^]. However, many of the most significantly differentially regulated genes in our data set were involved in cellular metabolism. In particular, 2 to 10 fold lower levels of arginine, serine, aspartate and glutamate led to a significant upregulation of rate-limiting steps in these pathways (*ASS1, PHGDH, PYCR1, GOT1*). Arginine levels in particular have been highlighted as critical for T cell proliferation, differentiation and survival (Rodriguez, Quiceno and Ochoa, 2007; Geiger *et al.*, 2016). The concentration of arginine in basal RPMI is roughly tenfold higher than in human plasma (1.2 mM versus 0.1 mM), and previous studies showing improved T cell survival with arginine supplementation use a concentration of 3 mM. Thus, it is likely that in the *in vivo* environment arginine is even more limiting than previously thought.

Broadly speaking, our data underscore the fact that commonly used cell culture conditions for T lymphocytes may not be ideal for metabolic studies and identifies one component in particular which can easily and rapidly be remedied by researchers in the field. This approach highlights the value of studying human physiologic media to improve the physiological relevance of *in vitro* techniques.

## ACKNOWLEDGEMENTS

M.L.G. was a student in the NIH-University of Pennsylvania graduate partnership program and the authors would like to acknowledge the mentorship and guidance provided. We thank Ann Park, Xijin Xu, other members of the Molecular Development of the Immune System Section at the NIH, John Wherry, Sara Cherry, Pamela Schwartzberg and Igor Brodsky for helpful discussions as well as technical guidance. We are grateful for financial support from the Lupus Foundation of America. The authors would also like to thank the Clinical Cancer Research Sequencing Facility for assistance in performing the RNA-Sequencing experiments and subsequent analysis, in particular; Justin Lack, Jyoti Shetty and Bao Tran. Lastly, the authors thank Tori Yamamoto for the kind gift of the CAR-T cell lentivirus. J.R.C. is supported in part by the NIH/NCI (K22CA225864). This research was supported in part by the Intramural Research Program of the NIH, National Institute of Allergy and Infectious Diseases.

## AUTHOR CONTRIBUTIONS

M.L.G., H.C.S. and M.J.L. initiated the project and designed the research. M.L.G. performed the experiments and analyzed the data with input from J.R.C., H.C.S. and M.J.L. J.R.C. prepared the HPLM media. A.B. performed bioinformatic analysis. M.L.G., H.C.S. and M.L. wrote the manuscript and all authors edited.

## MATERIALS AND METHODS

### Study Subjects

Anonymized blood samples were received from the NIH department of laboratory medicine and processed in order to isolate PBMCs.

### Cell Culture

In order to maintain consistency and reduce potential batch effects RPMI was prepared from powder (US Biologicals). Fresh L-glutamine (Gibco) and dialyzed FBS (Thermo Fisher Scientific) were added immediately prior to use at a final concentration of 2 mM and 10% by volume respectively. HPLM was prepared as previously described (Cantor *et al.*, 2017) along with four additional compounds: Uridine (3 µM), α-ketoglutarate (5 µM), acetylcarnitine (5 µM) and malate (5 µM). HPLM was supplemented with 10% dialyzed FBS throughout this study. HPLM-Min was prepared as HPLM for concentrations of ions, glucose and amino acids but polar metabolites were omitted (see Supplemental Table 1). RPMI was supplemented with 10% of either dialyzed (RPMI^dFBS^) or traditional FBS (RPMI^cFBS^). For long-term T cell proliferation assays, human recombinant IL-2 (Roche) was also added fresh immediately prior to use at a final concentration of 100 IU/ml. X-VIVO 15 media (Lonza) was used without additional serum. AIM-V media was a mixture of 50% RPMI and 50% AIM-V (Gibco) supplemented with 5% Normal Human Serum (Valley Biomedical). RPMI-Metabolites was prepared by adding the 31 small metabolites present in HPLM to basal RPMI and then supplementing with dialyzed FBS.

### Flow Cytometry

For simple surface stains, cells were washed once in phosphate buffered saline (PBS) and stained with 50 μl of Zombie Aqua™ viability (Biolegend) dye for 20 minutes on ice. After washing once with FACS buffer (PBS containing 5% FBS and 0.1% NaN_3_), cells were stained in 50 μl of FACS buffer containing diluted antibodies for 30 minutes on ice. Cells were then washed three times in FACS buffer and fixed before acquisition on either a LSRII or LSRFortessa (BD Biosciences). For intracellular cytokine staining following restimulation, cells were washed with PBS, stained in antibodies in 50 ul of FACS buffer for 30 minutes on ice. Cells were then washed three times in FACS buffer, then incubated for 20 minutes in 50 ul of Fixation/Permeabilization solution (BD Biolegend). Cells were washed three times in Perm/Wash buffer (BD Biolegend), then stained for 30 minutes on ice in antibodies diluted in Perm/Wash buffer. Cells were washed three times, then resuspended in Perm/Wash buffer and acquired on an LSRII. Data was analyzed using FlowJo v. 9.9.5.

### T cell Activation and expansion Analysis

Peripheral blood mononuclear cells were freshly isolated from blood from healthy human donors of both sexes ranging from 25-60 years old. Naïve T cells were enriched via a negative selection kit (Miltenyi Biotech, 130-097-095) and either used immediately or stored in liquid nitrogen in a 9:1 mixture of FBS: dimethyl sulfoxide. For short-term activation assays, 96 well Nunc Maxisorp plates were coated with 1 to 10 ug/ml of anti-CD3/anti-CD28 antibodies in PBS for 2 to 4 hours at 37°C. Plates were then washed twice with PBS and enriched naïve T cells were added in the indicated media formulations. Levels of activation markers were measured by flow cytometry 16 to 24 hours later. For proliferation assays, cells were stained with Celltrace Violet and then activated as above. After three days, cells were transferred to a new plate with fresh media and analyzed by flow cytometry after another two days.

### Calcium flux measurements

Purified naïve human T cells were cultured in RPMI for 16 to 24 hours following isolation or thawing from liquid nitrogen storage. Cells were then loaded with Indo-1 calcium Indicator dye (Thermo Fisher Scientific) resuspended in PowerLoad™ (Thermo Fisher Scientific) at a concentration for 20 minutes at room temperature. and stained for surface markers. Anti-CD3 (Hit3α) was added at 10 ug/ml, samples were acquired for 20 seconds to determine background levels, then F(ab’)_2_ fragments (Jackson Immunoresearch) were added at 13 ug/ml in order to crosslink the TCR. The flux was then recorded for another 130-180 seconds and analyzed by measuring the ratio of fluorescence in the two channels.

### T cell cytokine production assays

Human T lymphocytes were activated and expanded in IL-2-containing media as described above for 15 to 20 days. The night before the assay, cells were placed in fresh media (ether HPLM or RPMI) containing IL-2, and the following day were restimulated with either 1 μg/ml each of anti-CD3 (HIT3α, Biolegend) or PMA and ionomycin (Biolegend) for 6 or 5 hours respectively. Four hours prior to analysis, Brefeldin A was added (Biolegend). Following the restimulation, cells were washed in PBS, placed on ice and stained according to the protocol described above.

### Retroviral Transductions

CD19-CAR expressing retrovirus was produced as previously described (Kerkar *et al.*, 2011). 96-well plates were coated with Retronectin (Clontech) overnight at 4°C, washed and blocked with 2.5% bovine serum albumin (BSA), washed again and then bound to retrovirus by centrifuging the plates with viral supernatant for two hours at 37°C. The supernatants were then aspirated and human T lymphocytes that had been activated the previous day were added to the plate in the medium indicated. 48 hours following transduction, the cells were stained with Biotin-Protein L (Genscript) followed by fluorescently-conjugated streptavidin.

### RNA-Sequencing

RNA was isolated from cells using trizol (Thermo Fisher). We then used 0.1 – 1 ug of total RNA as input for mRNA capture with oligo-dT coated magnetic beads using the Illumina TruSeq protocol. The mRNA was fragmented, and then a random-primed cDNA synthesis was performed. The resulting double-strand cDNA was used as the input for a standard Illumina library prep with end-repair, adapter ligation and PCR amplification and then quantitated by qPCR followed by cluster generation and sequencing. RNA-Seq processing was conducting using the Pipeliner RNA-Seq workflow for quality assessment. For gene expression analysis, reads were trimmed to remove adapters and low-quality regions using Trimmomatic v0.33. Trimmed reads were aligned to the human GRCh38 reference genome and Gencode release 28 annotation using STAR v2.5.3 run in 2-pass mode. MultiQC v1.1 was used to aggregate QC metrics from picard, FastQC v0.11.5, FastQ screen v0.9.3 and RseQC to assess read and alignment quality. RSEM v1.3.0 was used for gene-level quantification and the resultant raw counts were voom-quantile normalized and batch corrected using the R package limma v3.38.3. Only genes that passed a 1 CPM threshold across a minimum of 3 samples based on the size of the smallest library were carried forward for differential expression testing. Pre-ranked gene set enrichment analysis (GSEA) was carried out using Gene Set Enrichment Analysis tool (GSEA v3.0) and Kegg pathways from MSigDB. Heatmap figures for visualization of gene expression and normalized enrichment score (NES) were generated using the heatmap.2 function from the R package gplots v3.0.1.1 and Clustvis respectively.

### Quantification and Statistical Analyses

One-way ANOVA with Dunnett’s/Tukey’s multiple comparisons test and Student’s t test were performed using Prism software (GraphPad). Descriptions of sample size and particular tests used can be found in the figure legend.

---

**Supplemental Figure 1 – GSEA of purified naïve human T cells activated in either HPLM or RPMI.**

Plot showing KEGG pathways enriched at either 48 or 120 hours following stimulation in either HPLM or RPMI.

