## Supplementary figures and images for "Human plasma-like medium improves T lymphocyte activation"

### Supplemental Figure 1

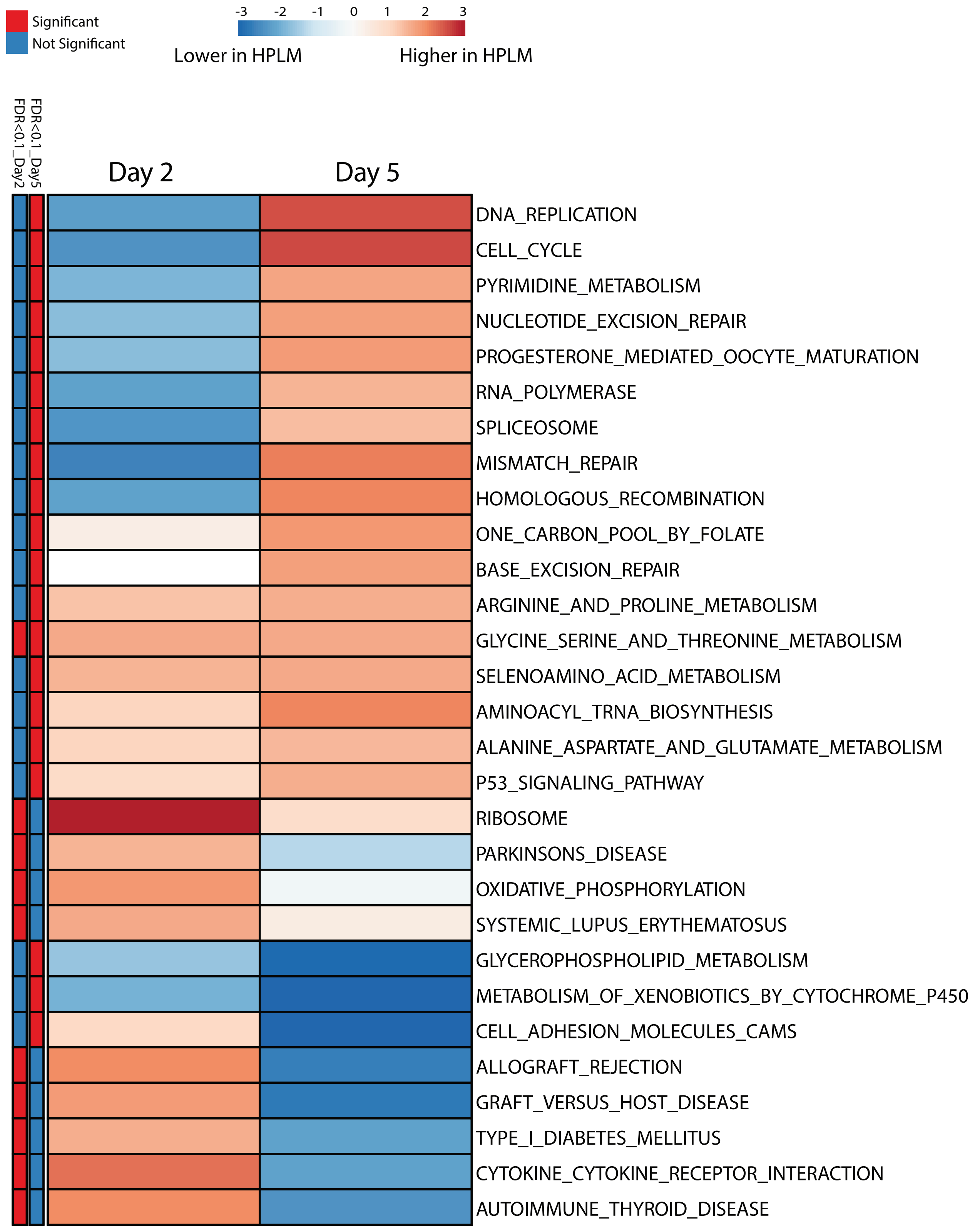


#### Supplemental Figure 1
