## Supplemental Table 1 for "Human plasma-like medium improves T lymphocyte activation"

### Supplemental Table 1: Detailed Media Composition

| | Concentration (μM) | | |
|---|---|---|---|
| | RPMI | HPLM-MIN | HPLM |
| **Glucose** | 11111 | 5000 | 5000 |
| ***Proteinogenic Amino acids*** | | | |
| **Alanine** | 0 | 430 | 430 |
| **Arginine** | 1149 | 110 | 110 |
| **Asparagine** | 378 | 50 | 50 |
| **Aspartate** | 150 | 20 | 20 |
| **Cysteine** | 0 | 40 | 40 |
| **Cystine** | 208 | 100 | 100 |
| **Glutamate** | 136 | 80 | 80 |
| **Glutamine** | 2055 | 550 | 550 |
| **Glycine** | 133 | 300 | 300 |
| **Histidine** | 97 | 110 | 110 |
| **Hydroxyproline** | 153 | 0 | 0 |
| **Isoleucine** | 382 | 70 | 70 |
| **Leucine** | 382 | 160 | 160 |
| **Lysine** | 219 | 200 | 200 |
| **Methionine** | 101 | 30 | 30 |
| **Phenylalanine** | 91 | 80 | 80 |
| **Proline** | 174 | 200 | 200 |
| **Serine** | 286 | 150 | 150 |
| **Threonine** | 168 | 140 | 140 |
| **Tryptophan** | 25 | 60 | 60 |
| **Tyrosine** | 11 | 80 | 80 |
| **Valine** | 171 | 220 | 220 |
| ***Ions*** | | | |
| **$Na^+$** | 138525 | 132271 | 132271 |
| **$K^+$** | 5333 | 4142 | 4142 |
| **$Ca^{2+}$** | 424 | 2390 | 2390 |
| **$Mg^{2+}$** | 407 | 830 | 830 |
| **$NH^{4+}$** | 0 | 40 | 40 |
| **$Cl^-$** | 108781 | 116196 | 116196 |
| **$HCO^{3-}$** | 23809 | 24000 | 24000 |
| **$PO_4^{3-}$** | 5634 | 966 | 966 |
| **$SO_4^{2-}$** | 407 | 350 | 350 |
| **$NO^{3-}$** | 848 | 80 | 80 |
| ***Additional Polar Metabolites*** | | | |
| **2-hydroxybutyrate** | | | 50 |

| | | | |
|---|---|---|---|
| **3-hydroxybutyrate** | | | 50 |
| **4-hydroxyproline** | | | 20 |
| **Acetate** | | | 40 |
| **Acetone** | | | 60 |
| **Acetylcarnitine** | | | 5 |
| **Acetylglycine** | | | 90 |
| **α-aminobutyrate** | | | 20 |
| **α-ketoglutarate** | | | 5 |
| **Betaine** | | | 70 |
| **Carnitine** | | | 40 |
| **Citrate** | | | 130 |
| **Citrulline** | | | 40 |
| **Creatine** | | | 40 |
| **Creatinine** | | | 75 |
| **Formate** | | | 50 |
| **Fructose** | | | 40 |
| **Galactose** | | | 60 |
| **Glutathione** | | | 25 |
| **Glycerol** | | | 120 |
| **Hypoxanthine** | | | 10 |
| **Lactate** | | | 1600 |
| **Malate** | | | 5 |
| **Malonate** | | | 10 |
| **Ornithine** | | | 70 |
| **Pyruvate** | | | 50 |
| **Succinate** | | | 20 |
| **Taurine** | | | 90 |
| **Urea** | | | 5000 |
| **Uric Acid** | | | 350 |
| **Uridine** | | | 3 |
